# The Shared Genetic Basis of Educational Attainment and Cerebral Cortical Morphology

**DOI:** 10.1101/242776

**Authors:** Tian Ge, Chia-Yen Chen, Alysa E. Doyle, Richard Vettermann, Lauri J. Tuominen, Daphne J. Holt, Mert R. Sabuncu, Jordan W. Smoller

**Affiliations:** Psychiatric and Neurodevelopmental Genetics Unit, Center for Genomic Medicine, Massachusetts General Hospital, Boston, MA 02114, USA; Department of Psychiatry, Massachusetts General Hospital, Harvard Medical School, Boston, MA 02114, USA; Athinoula A. Martinos Center for Biomedical Imaging, Massachusetts General Hospital, Charlestown, MA 02129, USA; Stanley Center for Psychiatric Research, Broad Institute of MIT and Harvard, Cambridge, MA 02138, USA; Analytic and Translational Genetics Unit, Center for Genomic Medicine, Massachusetts General Hospital, Boston, MA 02114, USA; School of Electrical and Computer Engineering and Nancy E. and Peter C. Meinig School of Biomedical Engineering, Cornell University, Ithaca, NY 14853, USA.

**Author notes:** Correspondence to Tian Ge.

## Abstract

Individual differences in educational attainment are linked to differences in intelligence, and predict important social, economic and health outcomes. Previous studies have found common genetic factors that influence educational achievement, cognitive performance and total brain volume (i.e., brain size). Here, in a large sample of participants from the UK Biobank, we investigate the shared genetic basis between educational attainment and fine-grained cerebral cortical morphological features, and associate this genetic variation with a related aspect of cognitive ability. Importantly, we execute novel statistical methods that enable high-dimensional genetic correlation analysis, and compute high-resolution surface maps for the genetic correlations between educational attainment and vertex-wise morphological measurements. We conduct secondary analyses, using the UK Biobank verbal-numerical reasoning score, to confirm that variation in educational attainment that is genetically correlated with cortical morphology is related to differences in cognitive performance. Our analyses reveal the genetic overlap between cognitive ability and cortical thickness measurements in bilateral primary motor cortex and predominantly left superior temporal cortex and proximal regions. These findings may contribute to our understanding of the neurobiology that connects genetic variation to individual differences in educational attainment and cognitive performance.

## Introduction

Educational attainment is a heritable trait [Krapohl et al., 2014; Polderman et al., 2015] that is predictive of many social, economic and health outcomes. Although linked to a range of diverse factors, it is highly phenotypically and genetically correlated with intelligence [Okbay et al., 2016; Rietveld et al., 2014; Savage et al., 2017; Sniekers et al., 2017], and has been successfully used as a proxy phenotype to facilitate the discovery of genetic variants associated with cognitive ability [Rietveld et al., 2014]. In addition, polygenic scores of educational attainment are predictive of cognitive performance in adolescents and adults [Belsky et al., 2016; Plomin and von Stumm, 2018; Selzam et al., 2017]. Dissecting the biological bases of educational achievement may thus contribute to our understanding of cognition and adult functional outcomes.

Large-scale genome-wide association studies (GWAS) have found substantial genetic overlap between educational attainment and total brain volume (i.e., brain size) [Adams et al., 2016; Okbay et al., 2016]. In particular, a recent GWAS of educational attainment has identified genomic loci regulating brain-specific gene expression, and biological pathways involved in neural development [Okbay et al., 2016]. Twin studies have implicated common genetic factors that influence both brain size and intelligence [Pol et al., 2006; Posthuma et al., 2002, 2003; Thompson et al., 2001; Toga and Thompson, 2005], a major contributor to the heritability of academic achievement [Krapohl et al., 2014]. Our recent investigation further suggested that genetic influences on individual differences in educational attainment might be mediated by brain development [Elliott et al., 2018]. However, to the best of our knowledge, no prior work has mapped the genetic correlations between educational attainment and fine-grained brain morphological measurements, likely due to methodological challenges and sample size (statistical power) limitations. Filling this knowledge gap represents an important next step in identifying the specific brain regions that lie in the pathway connecting genetics to educational outcomes.

In this study, we leverage the structural brain magnetic resonance imaging (MRI) scans and genomic data from a large sample of the UK Biobank participants (http://www.ukbiobank.ac.uk) [Sudlow et al., 2015] to investigate the shared genetic basis between educational attainment (years of schooling completed) and vertex-wise cortical thickness and surface area measurements. We also conduct secondary analyses, using the UK Biobank verbal-numerical reasoning score, to confirm that variation in years of education that is genetically correlated with cortical morphology is related to individual differences in cognitive ability. The verbal-numerical reasoning score assesses general cognitive ability and is heavily weighted towards math reasoning and vocabulary. Thus, it emphasizes learned knowledge, which has a strong relationship to educational attainment [Kaufman et al., 2009].

Well-established genetic correlation estimation methods such as genome-wide complex trait analysis (GCTA; also known as the GREML method) [Yang et al., 2011] and LD (linkage disequilibrium) score regression [Bulik-Sullivan et al., 2015a] require either individual genotypes or GWAS summary statistics for the two traits of interest. However, to examine the genetic overlap between educational attainment/cognitive performance (verbal-numerical reasoning) and massive numbers of brain morphological measurements in a large sample, both GCTA and LD score regression can be computationally intractable. For example, one would have to run thousands of GWAS in order to use LD score regression, and multiple testing correction would be challenging in the presence of complex spatial correlation structures. Here we develop a computationally efficient method that enables high-dimensional genetic correlation estimation, and the empirical identification of brain regions that are genetically correlated with our constructs of interest above and beyond global brain volumetric measurements. The power requirements for this method as well as our interest in relating genetic variation, cognition and real world functional outcomes dictate our approach. We first conduct genetic correlation analyses based on our larger sample with data on educational attainment, and then relate the variation in educational attainment that is genetically correlated with cortical morphology to differences in cognitive performance. These analyses expand the literature on the genetic underpinnings and brain morphological correlates of educational attainment, and may contribute to our understanding of the neurobiology of cognitive ability.

## Methods

### The UK Biobank

UK Biobank is a prospective cohort study of 500,000 individuals recruited across Great Britain during 2006-2010 [Sudlow et al., 2015]. The protocol and consent were approved by the UK Biobank’s Research Ethics Committee. Details about the UK Biobank project are provided at http://www.ukbiobank.ac.uk. Data for the current analyses were obtained under an approved data request (ref: 32568; previously 13905).

### Genetic data

The genetic data for the UK Biobank comprised 488,377 samples. Two closely related Affymetrix arrays were used to genotype ~800,000 markers spanning the genome. In addition, the dataset was phased and imputed to ~96 million variants with the Haplotype Reference Consortium (HRC) [Consortium, 2016] and the UK10K haplotype resource. We constrained all analyses to the HRC panel in the present study, which combines whole-genome sequence data from multiple cohorts of predominantly European ancestry, and thus covers a large majority of the common genetic variants in the European population.

The genetic data was quality controlled (QC) by the UK Biobank. Important information such as population structure and relatedness has been released. Details about the QC procedures can be found in Bycroft et al. [2017]. We leveraged the QC metrics provided by the UK Biobank and removed samples that had mismatch between genetically inferred sex and self-reported sex, high genotype missingness or extreme heterozygosity, sex chromosome aneuploidy, and samples that were excluded from kinship inference and autosomal phasing. We removed one individual from each pair of the samples that were 3rd degree or more closely related relatives, and restricted our analysis to participants that were estimated to have white British ancestry using principal component analysis (PCA).

### Brain imaging

We used the T1 structural brain MRI scans from 10,102 participants released by the UK Biobank in February 2017. FreeSurfer [Fischl, 2012] version 6.0 was used to process the MRI scans. All processed images were manually inspected and those with processing errors, motion artifacts, poor resolution, pathologies (e.g., tumors) and other abnormalities were removed. Among the 9,229 participants that passed imaging QC, a subset of 7,818 unrelated white British participants additionally passed the genetic QC described above and were included in the analysis. We resampled subject-specific morphological measurements (cortical thickness and surface area) onto FreeSurfer’s *fsaverage* representation, which consists of 163,842 vertices per hemisphere with an inter-vertex distance of approximately 1-mm. We further smoothed the co-registered surface maps using a surface-based Gaussian kernel with 20-mm full width at half maximum (FWHM).

### Educational Attainment

Following Okbay et al. [2016], we mapped each of the educational qualifications collected from the UK Biobank participants (UK Biobank field ID: 6138) to one of the seven categories defined in the 1997 International Standard Classification of Education (ISCED) of the United Nations Educational, Scientific and Cultural Organization, and imputed the number of years of schooling completed for each ISCED category. The mapping is shown in Supplementary Tables S1 and S2. Of all the participants that passed genetic QC, 332,613 (age, 39–72 y; female, 53.77%; years of education, 14.8±5.1 y) had years of schooling imputed at the baseline assessment visit (2006–2010) and were used in the GWAS of educational attainment. There was no overlap between the GWAS sample and the neuroimaging sample.

### Test of verbal-numerical reasoning

The verbal-numerical reasoning score (UK Biobank field ID: 20016; labeled as fluid intelligence score) used in the present study is an unweighted sum of the number of correct answers given to the thirteen higher-order reasoning questions [Lyall et al., 2016] in the UK Biobank touchscreen questionnaire. Participants who did not answer all of the thirteen questions within the allotted two-minute limit were scored as zero for each of the unattempted questions. Of all the participants that passed genetic QC, 108,147 (age, 40-70 y; female, 53.51%; verbal-numerical reasoning score, 6.2±2.1) had verbal-numerical reasoning scores at the baseline assessment visit (2006–2010) and were used in the GWAS. There was no overlap between the GWAS sample of the verbal-numerical reasoning score and the neuroimaging sample.

### GWAS of educational attainment and the verbal-numerical reasoning score

We performed GWAS of educational attainment (years of schooling completed) and the verbal-numerical reasoning score in 332,613 and 108,147 UK Biobank participants, respectively. In addition to the sample QC described above, we filtered out genetic markers with minor allele frequency < 1% and imputation quality score < 0.8. A total of 7,656,609 and 7,658,275 imputed SNPs on the HRC panel were included in the two GWAS, respectively. Association tests were conducted using SNPTEST v2.5.2 [Marchini and Howie, 2010]. For each genetic marker, a linear regression model was fitted, adjusting for age (at the baseline assessment visit), sex, age^2^, age× sex, age^2^ × sex, genotype array, UK Biobank assessment center, and top 10 principal components (PC) of the genotype data as covariates. GWAS results were visualized using FUMA [Watanabe et al., 2017] and the R package qqman [Turner, 2014].

### Estimators for SNP heritability

Consider the linear model ***y*** = ***X**β* + ***ϵ***, where ***y*** is an *N* × 1 vector of covariate-adjusted and standardized phenotypes, ***X*** = [*x_ij_*]*_N×M_* is an *N* × *M* matrix of genotypes with each column ***x**_j_* normalized to mean zero and variance one, *β* is an *M* × 1 vector of (random) SNP effect sizes, and *ϵ* is an *N* × 1 vector of residuals. In *Supplementary Information*, we show that under a polygenic model, the following moment-matching estimators for SNP heritability are asymptotically equivalent:

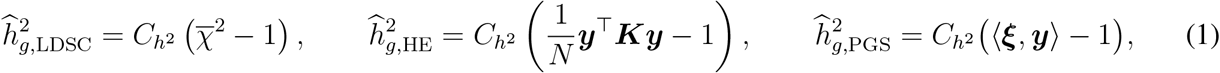

where 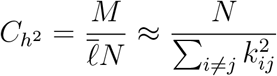 is a constant, 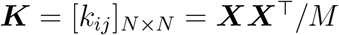 is the empirical genetic relationship matrix, 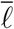 is the average **LD** score across the genome, 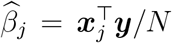 is the marginal effect size estimate of the *j*-th variant with 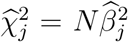 being the corresponding *χ*^2^ statistic, 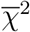 is the average *χ*^2^ statistic across the genome, 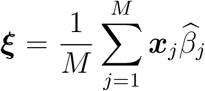 is a weighted average of the genotype, i.e., an *N* × 1 vector of individual-specific polygenic scores, and 〈·, ·〉 denotes inner product, i.e., 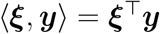.

We note that 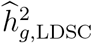 is the LD score regression estimator based on GWAS summary statistics, with the intercept constrained to one and the reciprocal of the LD score as the regression weight [Bulik-Sullivan et al., 2015b]. 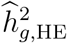 is the Haseman-Elston regression estimator based on individual genotypes [Elston et al., 2000; Ge et al., 2017a; Golan et al., 2014; Haseman and Elston, 1972]. 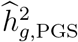 formulates SNP heritability estimation as a polygenic score analysis. The equivalence between 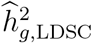 and 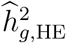 has been established both theoretically and empirically in prior work [Bulik-Sullivan, 2015; Ge et al., 2017a; Zhou, 2017].

### Estimators for SNP co-heritability

Consider the bivariate model ***y***_1_ = ***X***_1_*β*_1_ + *ϵ*_1_ and ***y***_2_ = ***X***_2_*β*_2_ + *ϵ*_2_, where ***y***_1_ and ***y***_2_ are *N*_1_ × 1 and *N*_2_ × 1 vectors of covariate-adjusted and standardized phenotypes, ***X***_1_ and ***X***_2_ are *N*_1_ × *M* and *N*_2_ × *M* matrices of standardized genotypes, *β*_1_ and *β*_2_ are *M* × 1 vectors of SNP effect sizes, *ϵ*_1_ and *ϵ*_2_ are *N*_1_ × 1 and *N*_2_ × 1 vectors of residuals, respectively. Without loss of generality, we assume that the first *N_s_* samples are identical for the two phenotypes. In *Supplementary Information*, we show that under a polygenic model, the following moment-matching estimators for SNP co-heritability are asymptotically equivalent:

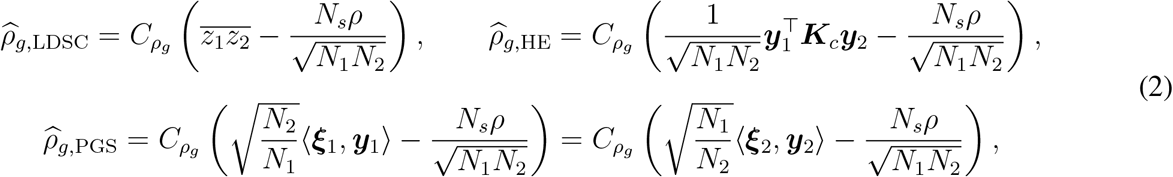

where 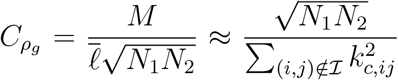 is a constant, 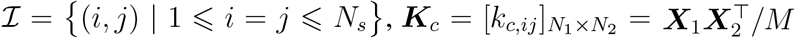, *ρ* is the phenotypic correlation between the two traits, 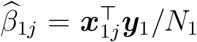 and 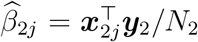 are marginal effect size estimates of the *j*-th variant, with 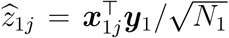 and 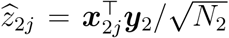 being the corresponding *z* statistics, respectively, 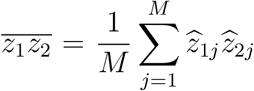 is the average product of *z* statistics across the genome, 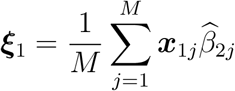 and 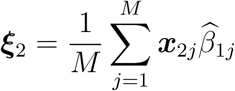 are individual-specific polygenic scores.

We note that 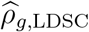 is the LD score regression estimator based on GWAS summary statistics, with a constrained intercept and the reciprocal of the LD score as the regression weight [Bulik-Sullivan et al., 2015a]. 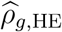 is the Haseman-Elston regression estimator based on individual genotypes. 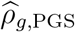 formulates SNP co-heritability estimation as a polygenic score analysis, and thus enables co-heritability analysis when GWAS summary statistics are available for one trait and individual genotypes are available for the other trait. The equivalence between 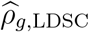 and 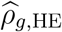 has been established in prior work [Bulik-Sullivan, 2015].

### Statistical genetic analyses

For all heritability and genetic correlation analyses, we used SNPs in the HapMap3 panel whose LD scores have been computed and released as part of the LD score regression software. We further filtered out genetic markers with imputation quality score < 0.9, missing rate > 1%, minor allele frequency < 1%, and significant deviation from Hardy-Weinberg equilibrium (*p* < 1 × 10^−10^) in the UK Biobank. A total of 870,962 SNPs were used in the heritability and genetic correlation analyses.

The SNP heritability of educational attainment, denoted as 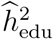, was computed using the LD score regression estimator 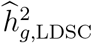 in Eq. (1) and the summary statistics of the education GWAS in the UK Biobank. The SNP heritability of the verbal-numerical reasoning score was computed similarly. The SNP heritability of the cortical thickness measurement at vertex *v*, denoted as 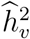, was computed using the Haseman-Elston regression estimator 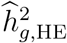 in Eq. (1) and individual genotypes of the imaging sample (*N* = 7, 818). We adjusted for age (at the imaging visit), sex, age^2^, age × sex, age^2^ × sex, handedness, genotype array, and top 10 PCs of the genotype data as covariates. We also controlled for the total brain volume (TBV) to remove global genetic influences on brain size. Vertex-wise estimates 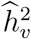, *v* = 1, 2, ⋯, *V*, where *V* is the total number of vertices, form a surface map for the heritability of cortical thickness measurements. The surface map for the heritability of surface area measurements was constructed similarly.

The SNP co-heritability between educational attainment and the cortical thickness measurement at vertex *v*, denoted as 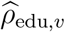, were computed using the estimator 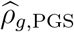 in Eq. (2). More specifically, the summary statistics of the education GWAS (*N* = 332,613) were used to calculate an individual-specific polygenic score in the imaging sample (*N* = 7,818) where individual genotypes were available. The polygenic score was then correlated with the cortical thickness measurement at each cortical location and properly scaled to produce the co-heritability estimate 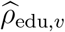. Since there was no overlap between the education GWAS sample and the neuroimaging sample, the bias term in the estimator, i.e., 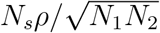, was set to zero. We adjusted for age (at the imaging visit), sex, age^2^, age × sex, age^2^ × sex, handedness, TBV, genotype array, and top 10 PCs of the genotype data as covariates in the co-heritability (polygenic score) analysis. The genetic correlation between educational attainment and the cortical thickness measurement at each vertex was then computed as

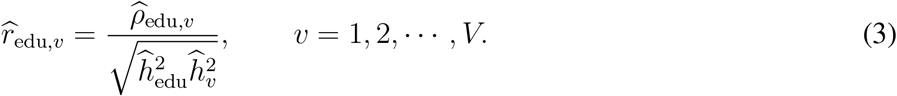

The genetic correlations between the verbal-numerical reasoning score and cortical thickness measurements were computed similarly.

Vertex-wise estimates 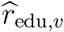, *v* = 1, 2, ⋯, *V*, form a surface map for the genetic correlations between educational attainment and cortical thickness measurements. Clusters on the surface map can be defined by spatially contiguous vertices with *p*-values below a threshold. To assess the significance of the size (number of vertices) of a cluster while accounting for the spatial correlation of cortical thickness measurements, we employed the following permutation procedure. For each permutation *k* = 1, 2, ⋯, *N*_perm_, we recomputed and thresholded the *p*-value map using a permuted polygenic score, and recorded the maximal cluster size *M_k_* across the two hemispheres. For an observed cluster 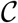 with size *c*, the family-wise error (FWE) corrected *p*-value was then computed as [Westfall and Young, 1993]

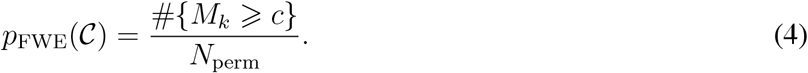

10,000 permutations were used in this study. We repeated the genetic correlation analyses using vertex-wise surface area measurements.

## Results

### GWAS of educational attainment

Genome-wide association analysis of educational attainment (years of schooling completed; *N* = 332, 613) in the UK Biobank identified 158 independent genome-wide significant loci. Figure 1A shows the Manhattan plot for the GWAS. Supplementary Figures S1 provides additional information on each of the genome-wide significant regions. The heritability of educational attainment was estimated to be 0.156 (s.e. 0.004).

**Figure 1:**
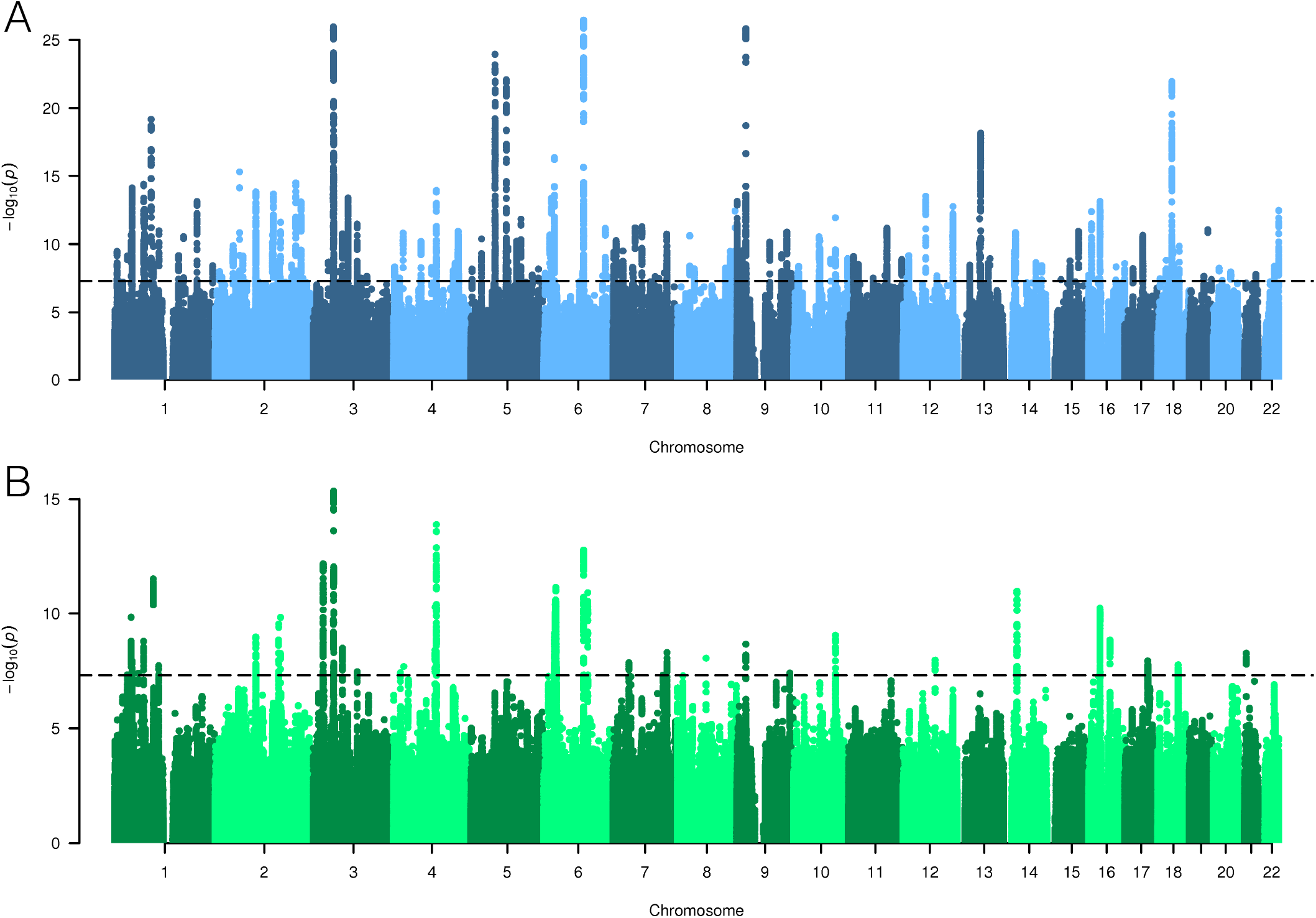
(A) Manhattan plot for the genome-wide association analysis of educational attainment (years of schooling completed) in the UK Biobank (*N* = 332, 613). (B) Manhattan plot for the genome-wide association analysis of the verbal-numerical reasoning score in the UK Biobank (*N* = 108,147). In both panels, the dash line indicates the genome-wide significant threshold *p* < 5 × 10^−8^.

### Heritability of cortical thickness

We estimated the SNP heritability of vertex-wise cortical thickness measurements using an unbiased and computationally efficient moment-matching method. As an empirical justification, our method produced virtually identical heritability estimates to LD score regression when applied to the average cortical thickness measurements in 68 regions of interest (ROIs; 34 ROIs per hemisphere) defined by the Desikan-Killiany atlas [Desikan et al., 2006] (Supplementary Figure S2, left; also see *Supplementary Information* for a theoretical treatment). As shown in Figure 2, fine-grained cortical thickness measurements were moderately heritable across the cortical mantle.

**Figure 2:**
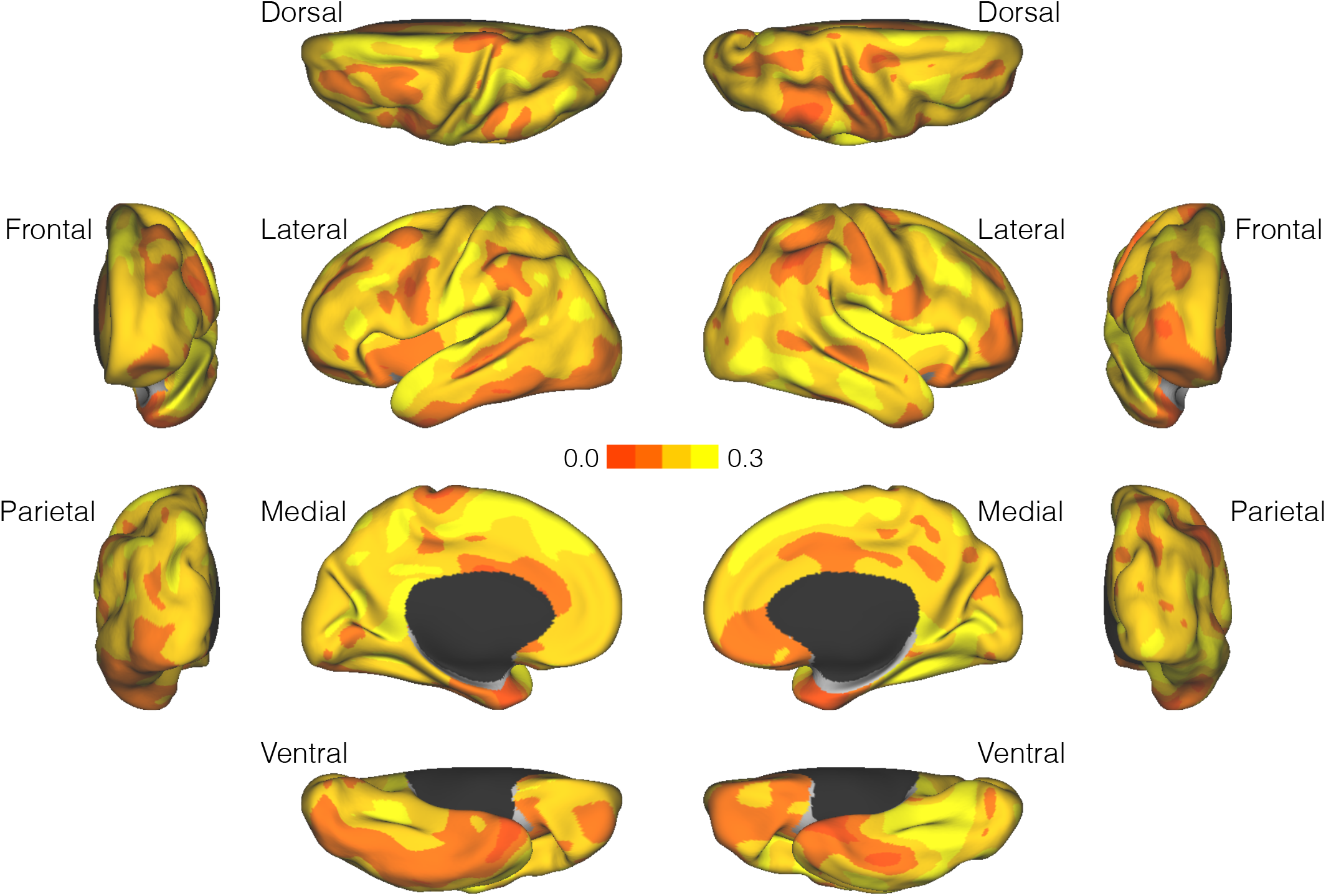
Surface maps for the SNP heritability of cortical thickness measurements (N = 7, 818).

### Genetic correlation between educational attainment and cortical thickness

Given that educational attainment and cortical thickness measurements were both heritable, we sought to examine whether they have a shared genetic basis. An empirical comparison of the genetic correlation between educational attainment and the average cortical thickness measurement in each of the 68 Desikan-Killiany ROIs estimated by the proposed polygenic score analysis and LD score regression showed that the two methods produced almost identical estimates (Supplementary Figure S2, right). Theoretical equivalence between the two methods is established in *Supplementary Information*.

Figure 3A and 3B show surface maps for the genetic correlation and its statistical significance between educational attainment and cortical thickness measurements, respectively. Moderate and positive genetic correlations were observed in bilateral motor cortex and predominantly left superior temporal cortex and proximal regions. We thresholded the significance map using p = 0.01 as the threshold (Figure 4B), and assessed the significance of the size of each identified cluster (spatially contiguous vertices) and computed their family-wise error (FWE) corrected *p*-values using a permutation procedure we devised. Statistically significant clusters were observed in bilateral primary motor cortex (cluster 1, *p*_FWE_ = 0.033; cluster 2, *p*_FWE_ = 0.042; and cluster 4, *p*_FWE_ = 0.012). Cluster 2 on the left hemisphere also extended into the pars opercularis (also known as Brodmann area 44 or BA44), which is part of the Broca’s speech area. In addition, cluster 3 (*p*_FWE_ = 0.005) spanned the left temporal pole and superior temporal cortex, and overlapped with the auditory cortex and the Wernicke’s language area. The right inferiortemporal cortex (cluster 5) also showed strong genetic correlation with educational attainment but did not survive multiple testing correction (*p*_FWE_ = 0.110).

**Figure 3:**
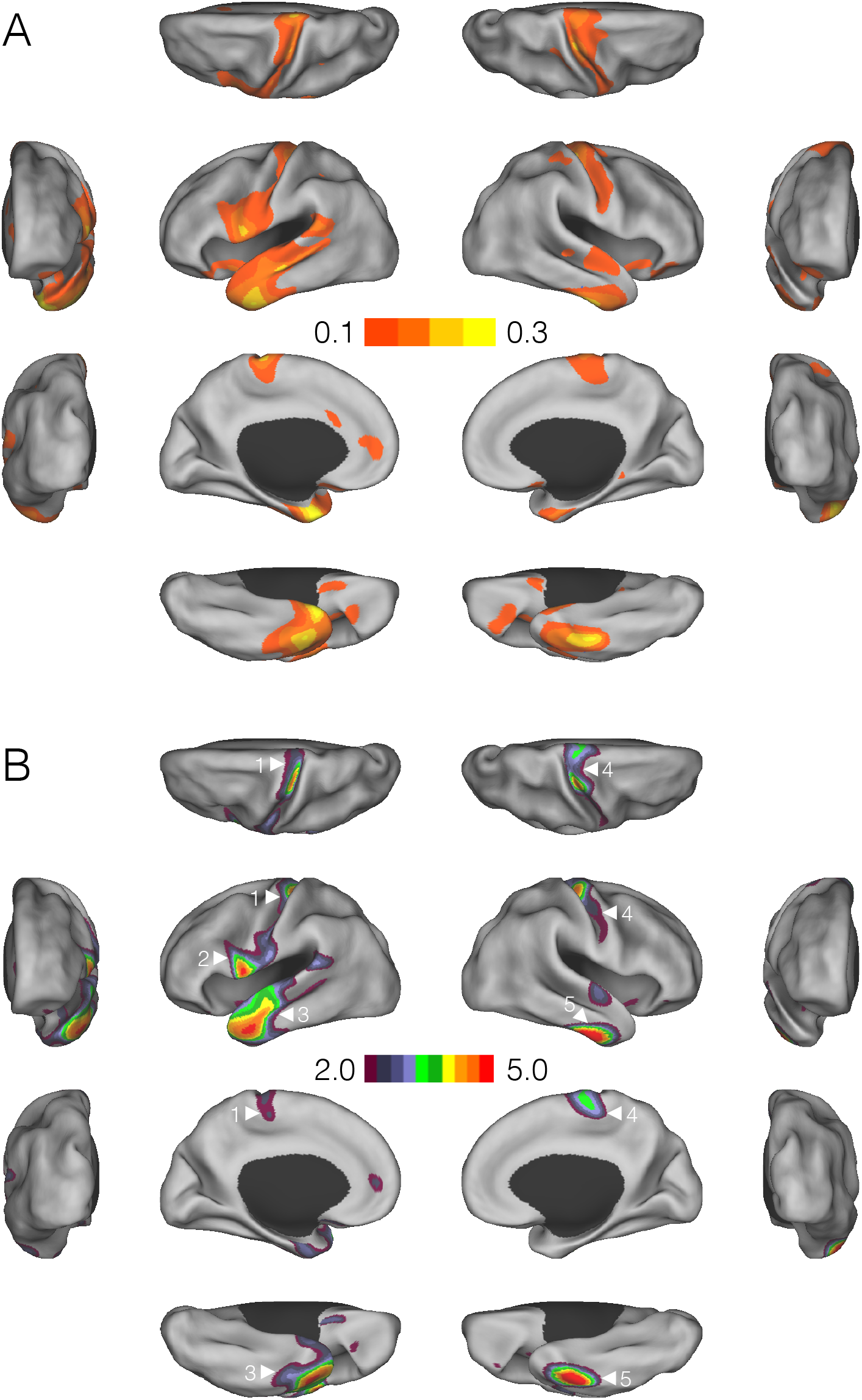
Vertex-wise genetic correlations between educational attainment and cortical thickness measurements. (A) Surface maps for the genetic correlation estimates. (B) Surface maps for the – log_10_ *p*-values of the genetic correlations. Clusters identified by a cluster-forming threshold of *p* = 0.01 are shown. Family-wise error corrected significant (or marginally significant) clusters are annotated.

**Figure 4:**
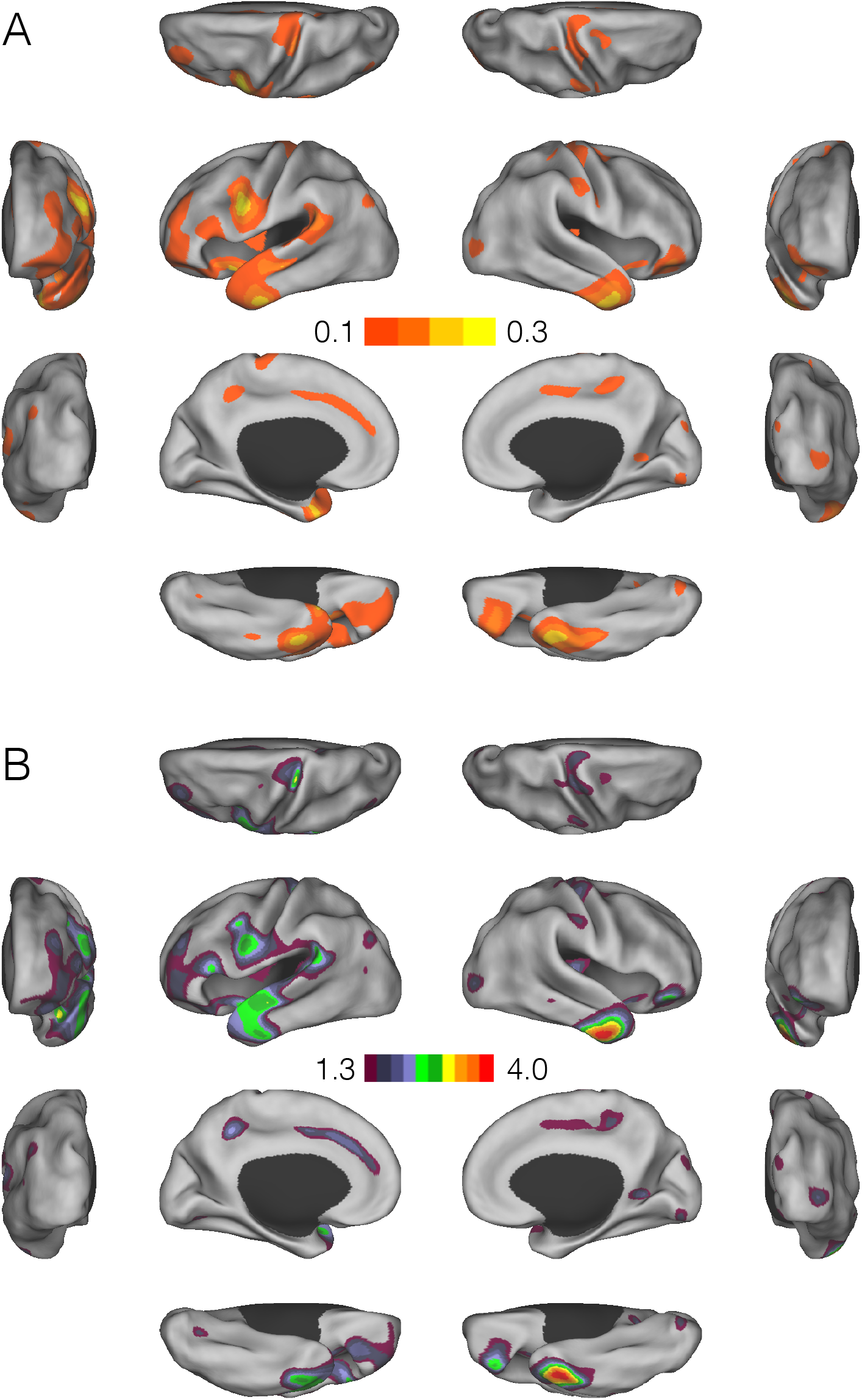
Vertex-wise genetic correlations between the verbal-numerical score and cortical thickness measurements. (A) Surface maps for the genetic correlation estimates. (B) Surface maps for the – log_10_ *p*-values of the genetic correlations, thresholded at uncorrected *p* = 0.05.

### Analyses of the verbal-numerical reasoning score

Analyses of educational attainment indicated that cortical thickness in several regions, including the primary motor cortex and Broca’s speech and Wernicke’s language areas, may have shared genetic origins with cognitive ability. We thus conducted a secondary analysis using the verbal-numerical reasoning score in the UK Biobank, which captures general cognitive ability but particularly knowledge of learned material, to investigate whether the variation in years of schooling that is genetically correlated with cortical thickness is related to individual differences in cognitive performance.

Genome-wide analysis of the verbal-numerical reasoning score in the UK Biobank (N = 108,147) identified 35 genome-wide significant loci. Figure 1B shows the Manhattan plot for the GWAS. Supplementary Figures S3 provides additional information on each of the genome-wide significant regions. The heritability of the verbal-numerical reasoning score was 0.247 (s.e. 0.008). Educational attainment and the verbal-numerical reasoning score were both phenotypically (Pearson correlation *r* = 0.353, *N* = 110, 233) and genetically (genetic correlation *r_g_* = 0.710; s.e. 0.016) correlated.

Figure 4A and 4B show surface maps for the genetic correlation and its statistical significance (thresholded at the nominal *p* = 0.05) between the verbal-numerical reasoning score and cortical thickness measurements, respectively. Both maps showed similar patterns to the surface maps for educational attainment, with positive genetic correlations observed in the predominantly left inferior precentral gyrus (including Broca’s speech area), superior temporal cortex (including auditory cortex), supramarginal gyrus (including Wernicke’s language area) and proximal regions. The genetic correlation estimates showed a similar range to those between educational attainment and cortical thickness measurements, but were less statistically significant due to the less powerful verbal-numerical reasoning GWAS relative to the education GWAS.

### Surface area analyses

We repeated all analyses using surface area measurements. Vertex-wise surface area measurements were substantially less heritable than cortical thickness measurements (Supplementary Figure S4). In contrast to cortical thickness, no significant cluster of genetic correlations between educational attainment and surface area measurements was identified (all *p*_FWE_ > 0.10; Supplementary Figure S5).

## Discussion

In this paper, we examined the genetic overlap between educational attainment and fine-grained brain morphological measurements. Leveraging the large-scale brain imaging and genomic data in the UK Biobank, we found a shared genetic basis between years of schooling and cortical thickness measurements in bilateral primary motor cortex and predominantly left superior temporal cortex. A secondary analysis of the verbal-numerical reasoning score confirmed that the variation underlying education achievement that is genetically correlated with cortical thickness is related to individual differences in cognitive performance.

Although educational attainment is a complex behavioral trait that is linked to intelligence, personality, family environments and many social factors, it is highly phenotypically and genetically correlated with intelligence [Okbay et al., 2016; Savage et al., 2017; Sniekers et al., 2017], and has been successfully used as a proxy phenotype to identify genetic variants associated with general cognitive ability [Rietveld et al., 2014]. Recent studies have associated the polygenic scores of educational attainment with cognitive test scores and brain size [Belsky et al., 2016; Plomin and von Stumm, 2018; Selzam et al., 2017], and further implicated that genetic influences on educational outcomes might be mediated through brain development and intelligence [Elliott et al., 2018]. Within the UK biobank, tests of cognitive functioning were brief and only completed by a sub-sample of the participants. In contrast, educational qualifications, which can be mapped directly to years of schooling, were available for virtually all UK Biobank participants. Therefore, we leveraged educational attainment as a proxy for general cognitive ability in the identification of genetic overlap between cerebral cortical morphology and cognitive performance. We used this trait in our discovery analysis because its size and objective reliability may boost the statistical power relative to the brief cognitive screening tests with substantially smaller sample sizes in the UK Biobank.

Of the thirteen questions that comprise the UK Biobank verbal-numerical reasoning score, six relate to numeric reasoning, four relate to vocab/verbal reasoning and three are logic questions. Conventionally, the ten verbal and numeric reasoning questions would fall within the domain of crystallized ability, which reflects learned knowledge, including material absorbed in the educational setting [Wilhelm, 2004]. Logic questions relate to novel problem-solving, and fall within the domain of fluid intelligence [Wilhelm, 2004]. Thus, our cognition measure, which is labeled as “fluid intelligence” in the UK Biobank directory, captures both crystallized and fluid intelligence (the major domains in a commonly accepted model of general ability) [Horn and Cattell, 1966], but is more heavily loaded towards crystallized knowledge. Given that crystallized ability would be expected to increase over time whereas fluid ability would decrease, our conceptualization of this measurement is consistent with Lyall et al. [2016] that this score in the UK Biobank improved slightly over a roughly four-year period. Measurements of crystallized ability are well-known to relate to educational attainment, and more recently measurements of fluid ability have been associated with this outcome [Kaufman et al., 2009]. As such, educational attainment is well-suited to serve as a proxy index of this cognitive ability measurement in our analyses.

Our analyses of educational attainment and the verbal-numerical reasoning score produced highly similar surface maps for genetic correlations and localized a common genetic basis between cognitive performance and cortical thickness measurements in bilateral primary motor cortex and predominately left inferior precentral gyrus, pars opercularis, superior temporal cortex, supramarginal gyrus, and their adjacent regions. Intriguingly, some of these regions overlap with the auditory cortex, Broca’s speech area and Wernicke’s language area, suggesting that educational attainment and our cognitive measurement may have common genetic origins with auditory and language-related brain regions. The distinctiveness of language and cognition has been debated over the 20th century [see e.g., Harris, 2003]. Our data provides evidence of potentially shared biological underpinnings that, if confirmed, could contribute to this line of inquiry. The primary motor cortex and its vicinity have been found to be consistently activated during reasoning tasks [Acuna et al., 2002; Prado et al., 2011], and implicated in lesion mapping of intelligence [Gläscher et al., 2010; Woolgar et al., 2010]. Therefore, our results also suggest that cognitive performance may have a shared genetic basis with brain regions involved in motor processes.

Previous studies have used neuroimaging to examine the relationships between intelligence, cognitive ability, education and brain structure *in vivo* [Cox et al., 2016; Luders et al., 2009; Sabuncu et al., 2016]. Higher intelligence has been associated with larger brains [McDaniel, 2005; Pietschnig et al., 2015; Rushton and Ankney, 2009; Wickett et al., 2000; Witelson et al., 2006], and greater total and regional gray matter volumes [Andreasen et al., 1993; Colom et al., 2006; Flashman et al., 1997; Haier et al., 2004, 2009; MacLullich et al., 2002]. Beyond total brain volume, positive correlations have been reported between scores of cognitive tests and cortical thickness measurements in a number of cortical regions [Choi et al., 2008; Joshi et al., 2011; Karama et al., 2009; Menary et al., 2013; Narr et al., 2007]. Our results expanded this literature by identifying brain regions in which cortical thickness measurements are genetically correlated with cognitive performance above and beyond global brain volumetric measurements. Cortical surface area has also been associated with general cognitive ability [Colom et al., 2013; Fjell et al., 2013; Schnack et al., 2014], and some recent studies have suggested that the phenotypic and genetic correlations between brain volume and cognitive performance are predominantly driven by surface area rather than cortical thickness [Cox et al., 2018; Vuoksimaa et al., 2014]; however, we did not observe significant genetic correlations between educational attainment and surface area measurements in our analyses.

A major contribution of this study is that we developed a novel statistical method to examine the shared genetic basis between educational attainment and cortical morphology at high spatial resolution. Prior studies along this line used the twin design and largely focused on global brain volumes. Existing methods for genetic correlation analyses in unrelated individuals either require individual-level genotypes (e.g., GCTA or GREML) [Yang et al., 2011] or GWAS summary statistics (e.g., LD score regression) [Bulik-Sullivan et al., 2015a] for both traits, which become challenging to apply in high-dimensional settings. We filled this technical gap by formulating genetic correlation estimation as a polygenic score analysis. More specifically, the summary statistics of the educational attainment (or verbal-numerical reasoning) GWAS were used to weight individual genotypes of the imaging sample and calculate an individual-specific polygenic score. The polygenic score was then correlated with the morphological measurement at each cortical location and properly scaled and normalized to produce a genetic correlation estimate. Our method is thus highly computationally efficient and can be applied to estimate the genetic correlation between any trait (e.g., a cognitive, behavioral or disease phenotype) whose GWAS summary statistics are available, and a high-dimensional phenotype, such as the MRI-derived vertex-wise cortical thickness or surface area measurements in the present study. Permutation procedures can also be devised to enable flexible statistical inferences, such as the cluster-wise analysis [Friston et al., 1994, 1996] on the surface map of genetic correlations. Standard polygenic score analyses may achieve similar results and findings but our method avoids the selection of the *p*-value threshold (or screening multiple thresholds) used in the calculation of polygenic scores, and produce more interpretable estimates (i.e., genetic correlation) at no additional computational cost.

To apply this new genetic correlation estimation method, we conducted genome-wide association analyses of educational attainment and the verbal-numerical reasoning score in the UK Biobank, and identified 158 and 35 independent genome-wide significant loci, respectively. Previous studies have conducted large-scale genome-wide meta-analyses of educational attainment [Okbay et al., 2016] and general intelligence [Savage et al., 2017; Sniekers et al., 2017], and GWAS with a similar or larger total sample size than the present study exists. Although we could have leveraged the summary statistics of existing GWAS in our analyses, we performed GWAS in the UK Biobank to exclude all participants that had neuroimaging data, and thus protected the genetic correlation estimates from potential bias induced by sample overlap.

Findings in this analysis should be generalized with caution to populations with different sample characteristics with respect to age range, sex composition, ancestry groups, socioeconomic status (SES), or other environmental exposures. Educational attainment in our analysis reflects life-time academic achievement, while both human intelligence and the cortex undergo rapid development in childhood and adolescence, and age-related decline and degeneration in late adulthood. More importantly, a number of studies have found that the relationship between cognitive ability and brain morphology is dynamic over time and sex-dependent [see e.g., Burgaleta et al., 2014; Gur et al., 1999; Reiss et al., 1996; Shaw et al., 2006; Witelson et al., 2006]. In this study, we controlled for age, sex, age^2^, age× sex, age^2^ × sex in all the analyses to remove the (potentially nonlinear) effect of age and sex on cognitive performance, brain structure, and their correlations. However, since the UK Biobank only recruited middle- and older-aged participants, our results may not be generalizable to other age ranges. In addition, genetic influences on cognitive performance may be moderated by educational opportunities and SES [Deary and Johnson, 2010; Hanscombe et al., 2012; Von Stumm and Plomin, 2015]. Therefore, our findings should be interpreted in light of the fact that UK Biobank participants are on average more educated and have higher SES than the general population [Fry et al., 2017; Tyrrell et al., 2016].

Although we identified genetic overlap between educational attainment and cortical thickness measurements in several brain regions, these correlations do not necessarily indicate causal relationships. Future work is needed to shed light on whether genetic influences on cognitive performance are mediated through brain morphology. Also, genetic correlation is a genome-wide metric and does not provide any information about specific genes that might underlie both cognitive ability and brain structure. Further statistical and molecular genetic analyses are needed to dissect their genetic overlap. Lastly, in addition to gray matter volumes, cortical thickness and surface area measurements, white matter volumes, diffusion tensor imaging (DTI) derived measurements, functional MRI task activations, and indices of complex brain networks have also been associated with cognitive performance [Deary et al., 2010; Luders et al., 2009]. Given that a range of features derived from brain morphology, resting state networks, and the structural and functional connectomes are substantially heritable [Ge et al., 2015, 2016, 2017b; Glahn et al., 2010; Kochunov et al., 2015; Thompson et al., 2013], integration of multimodal imaging data might provide further insights into possible neural mechanisms of educational attainment and cognitive ability.

## Acknowledgements

This research was carried out in part at the Athinoula A. Martinos Center for Biomedical Imaging at the Massachusetts General Hospital (MGH), using resources provided by the Center for Functional Neuroimaging Technologies, P41EB015896, a P41 Biotechnology Resource Grant supported by the National Institute of Biomedical Imaging and Bioengineering (NIBIB), National Institutes of Health (NIH). This work involved the use of instrumentation supported by the NIH Shared Instrumentation Grant Program (grant numbers S10RR023043 and S10RR023401), and the Enterprise Research Infrastructure & Services (ERIS) at Partners Healthcare. This research was also funded in part by NIH grants K99AG054573 (TG); R01MH095904 and R01MH109562 (DJH); R01AG053949 and R01LM012719 (MRS); and K24MH094614 (JWS). JWS is a Tepper Family MGH Research Scholar and was also supported in part by a gift from the Demarest Lloyd, Jr. Foundation. The funders had no role in study design, data collection and analysis, decision to publish, or preparation of the manuscript. This research has been conducted using the UK Biobank resource under an approved data request (ref: 32568; previously 13905).

